# 31° South: Dietary niche of an arid-zone endemic passerine

**DOI:** 10.1101/508465

**Authors:** Ângela M. Ribeiro, Ben Smit, M. Thomas P. Gilbert

## Abstract

Balancing energy budgets is thought to be challenging for birds living in arid ecosystems because food supplies are low and unpredictable, and climatic conditions extreme. Thus, to ensure they obtain sufficient energy to fuel daily energetic budgets, birds may need to adjust their diets and become less selective (generalist) as conditions become harsher. To test this hypothesis, we used DNA metabarcoding to characterize both the prey availability (from pitfall traps) and the dietary content (from fecal samples) of several conspecific populations of a semi- and arid-endemic insectivorous bird, the Karoo-scrub-robin (*Cercotrichas coryphaeus*) across a climatic gradient. Our results showed that Coleoptera, Hymenoptera, Orthoptera, and Lepidoptera were the main prey. When accounting for their presence as available prey, Coleoptera and Hymenoptera were preferred in all regions, whereas robins avoided Orthoptera and Lepidoptera in all but the most arid region. Although the different populations live in regions that vary with regards to productivity and thermoregulatory demands, we found that the dietary niche breadth (Bs) of the three populations was intermediate to low, and did not differ significantly. As a whole, our findings show that regardless of environmental harshness these insectivores have similar dietary niches, suggesting that large dietary plasticity is fundamental for their survival in energy-depauperated ecosystems.

## INTRODUCTION

Food acquisition is a basic need of all animals: it provides the energy and nutrients required to sustain life in any given environment. Because of the intricate relationship of animals with the environment (where food is obtained from), there has long been an interest in describing patterns of dietary variation in order to predict the underlying causes. The classical Optimal Foraging Theory model (MacArthur and Pianka 1966) (Levins and MacArthur 1969) (Schoener 1971) predicts that animals seek to maximize energy intake, so as to meet their daily energy costs. Consequently, both prey availability in the environment and animal physiological status can affect diet variation (e.g.: (Burgar et al. 2014).

Endotherms living in landscapes characterized by dramatic change in rainfall and temperature face two main problems: firstly, frequent changes in food webs; and secondly, thermal challenges that require them to adjust their physiology (energetically expensive tasks) to maintain body temperatures within tolerable limits. As a consequence of both, some populations may undergo dietary niche shifts in order to maximize fitness (Roches et al. 2016). In response to limited resources, selection can either lead to generalist strategies such as dietary niche flexibility to use the panoply of resources available (Grant et al. 2008), or drive the evolution of specialization so as to exploit underused resources (Martin and Pfennig 2009). Moreover, if resources vary greatly through time or space, being a generalist may allow fitness to be maintained (Stephens and Krebs 1986).

Arid-zone endemic birds are good models for testing the role of environmental temperature and energetic physiology in dietary strategies, because they face continual challenges from their energy-depauperated ecosystems, while seeking to fulfill the dietary requirements needed to fuel the physiological processes associated with thermoregulation. In this study we molecularly characterized both the diet and available arthropod prey community (Pompanon et al. 2012) (Bohmann et al. 2011) of conspecific populations of the Karoo-scrub robin (*Cercotrichas coryphaeus*; hereafter: robin), a ground-feeding insectivore (Roberts et al. 2005). This southern African passerine provides a good system with which to explore the avian dietary strategy to ensure adequate energy intake, because its range spans an environmental gradient of primary productivity variation (Fig. 1A). Primary productivity ultimately determines arthropod biomass in the regions, which in turn affects the food available for insectivores (Maclean 1969) (Lloyd 1999). Furthermore, these robins also face dramatic temperature changes, which lead thermoregulatory challenges that the birds must adjust their rates of energy expenditure to (Fig. 1B).

**Figure 1.**
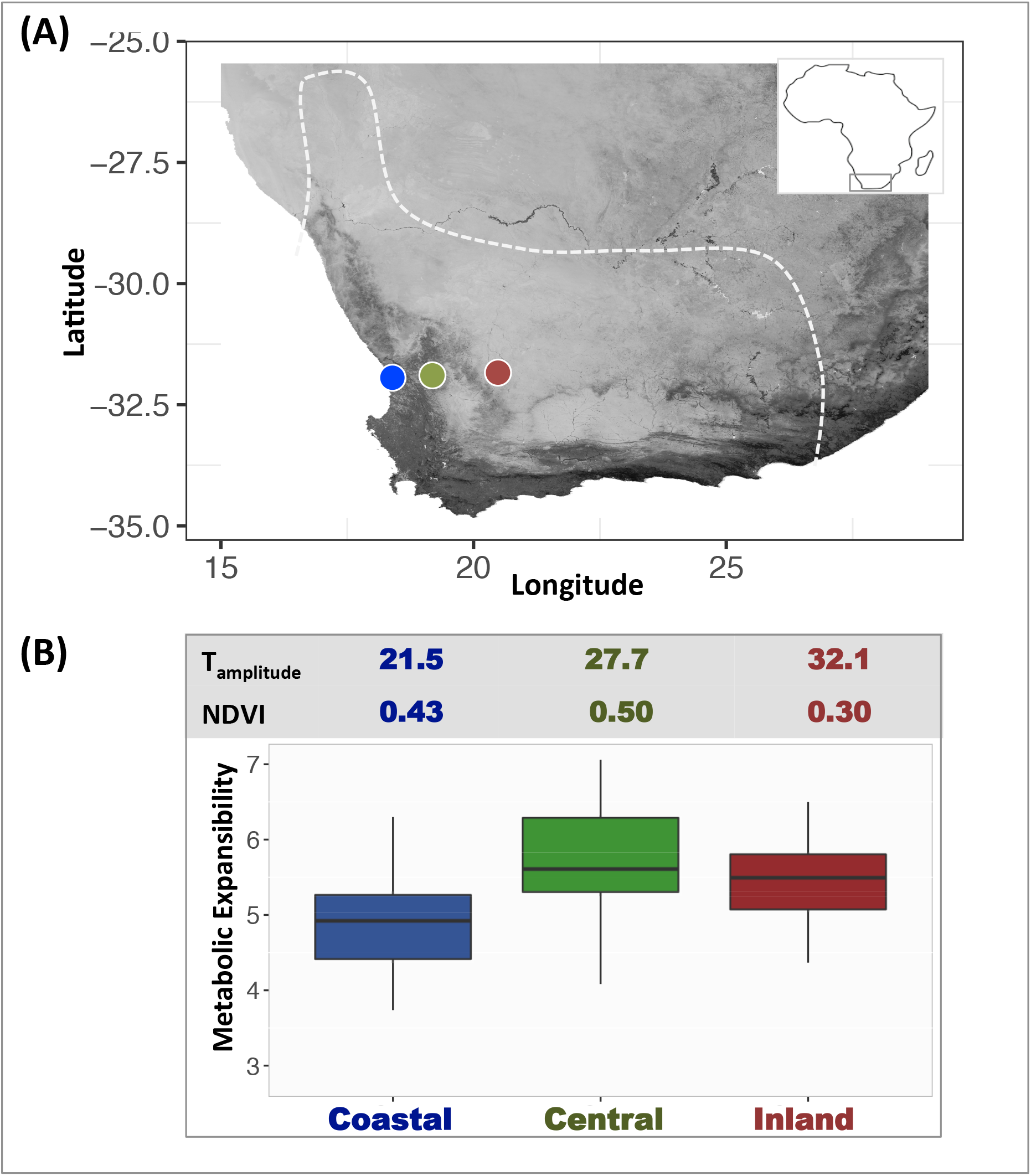
Energetic landscape of the three study sites and populations of Karoo scrub-robin (*Cercotrichas coryphaeus*). (A) Map of annual primary productivity (NDVI; the darker the pixel the larger the productivity) with depiction of the three study sites. Karoo scrub-robin range limit is delimited by a dashed line. (B) Primary productivity (NDVI), Temperature amplitude (°C) and energetic physiology (metabolic expansibility) for the study populations (from Ribeiro et al unpublished data) three study sites.

Our objectives were therefore two-fold. Firstly, to determine whether environmental heterogeneity resulted in differential prey availably. And secondly, to assess whether this prey availably and physiological status lead to differences in the birds’ niche breadths, a proxy for their dietary flexibility and prey selectivity. In particular, we hypothesized that populations resident in the area with the lowest productivity and highest metabolic expansibility (capacity to increase metabolism so as to deal with any energetic demand, for instance cold), should exhibit the most generalist feeding strategy, as their means for coping with the harshness of the arid-conditions.

## 2. MATERIAL AND METHODS

### 2.1 Study sites, arthropods and fecal sample collection

The study was conducted along a gradient of (i) annual temperature amplitude (T_amp_) and (ii) primary productivity (measured as normalized difference vegetation index – NDV) (Fig. 1). We selected these two metrics with the rationale that: wide ambient temperature range implies increased energy expenditure so as to deal with extreme climatic conditions (cold and hot), and consequent incremental intake of energy sources (Bicudo et al. 2010); and that NDVI is a good predictor of plant biomass in the arid-zone of southern Africa (Borer et al. 2012).

We used the GPS coordinates of our pitfall grid to obtain T_amp_ and NDVI. T_amp_ was estimated using data from WorldClim2 at 2.5min resolution (Fick and Hijmans 2017) as: maximum temperature in December – minimum temperature in July. NDVI data, the surrogate for primary productivity, was extracted from eMODIS dataset hosted by NASA-LPDAAC (USGS, USA). Beside environmental variables, we gathered physiological phenotypic data (Ribeiro et al. *in prep*) for populations inhabiting the same localities. We used these metrics to define three operational regions: *Coastal, Central,* and *Inland.*

We sampled available prey arthropods using 27 pitfall traps left open for seven consecutive days, at each of three locations (Fig. 1) in the summer of 2015 and winter of 2016. We selected pitfall traps because scrub-robins are mostly ground-feeders. The pitfall traps were placed within an area we previously checked to be used by robins, and then set them in three grids. Each grid comprised of nine traps forming a square (3m x 3m), with a distance of one meter between individual traps, and a distance of 100 m between grids. The traps were made from two nested transparent plastic cups of 8 cm diameter, and 250 ml of volume; we partially filled them with a solution of water and soap to reduce superficial tension (Cheli and Corley 2010). We collected the arthropods in the traps on the fourth and seventh (last) day of sampling. All arthropods caught per site were pooled to represent the arthropod community of the region in the given season. Post sampling, arthropods were washed with water and ethanol to remove debris, and stored in 96% ethanol until further use.

In the same above-mentioned periods we collected 49 bird fecal pellets (N_Coastal_= 16, N_Central_= 16, N_Inland_= 17) using single use filter-paper bags lining cloth bags in which captured birds were placed. Birds were caught using baited flat traps, in place from early morning to dusk in all three regions. Upon collection, the fecal pellets were placed in tubes containing beads and lysis buffer as provided in the Soil/Fecal DNA MiniPrep kit (ZymoResearch, USA), homogenized by vortexing at 2700rpm (45HZ) for 1 min, and stored at 4°C until further processing in the lab. The sampling periods were selected to avoid the breeding season; in fact, we found no sign of breeding activity while in the field.

### 2.2 DNA extraction

Prior to DNA extraction, arthropods collected from pitfall traps were sorted by size into three categories: small (<1cm), medium (1-3cm) and large (>3cm). The rational was two fold: first, large-sized arthropods could be easily identified morphologically (Appendix Table S1); and second, we wanted to extract DNA from several arthropods at a time (mixed sample), hence it is crucial that the animals in the same pool have similar sizes so not to biases the proportion of DNA in the final volume. Arthropod samples were drained and let dried at 56°C to remove all ethanol. In order to maintain the arthropod exoskeletons intact, we extracted DNA from arthropod samples following (Gilbert et al. 2007) and (Nielsen 2015). Briefly, this consisted of immersing the arthropods in a digestion buffer with incubation overnight at 56°C with gentle agitation, after which the digest was purified using the QiaQuick Purification kit (Qiagen, USA). Extracted DNA was stored at −20°C until further use.

We homogenized fecal samples prior to DNA extraction using a tissue lyser (Qiagen Tissue Lyser II, USA) for 30s at 30Hz, followed by 30s at 20Hz. DNA extractions used the Soil/Fecal DNA MiniPrep kit (ZymoResearch, USA), with one minor change to the manufacturer’s guidelines: after homogenization, we added Proteinase K and incubated the samples at 60 °C for 30 min. The DNA extracted was stored at −20°C until further use. We specifically chose this kit because it contains beads in a bead beating tube, a feature that may help crush both the exoskeleton fragments of arthropods and seeds in fecal pellets, hence increase the yield of DNA. Negative controls were included in each extraction to check for potential contamination. These negative controls proceeded in the workflow (from PCR to sequencing) as with all other extracts.

### 2.3 Metabarcoding and sequencing

To maximize detection and identification of arthropods we used two different primers targeting different portions of the COI gene in the mitochondrial genome of arthropods: i) ZBJ-ArtF1c/ZBJ-ArtR2c (Zeale et al. 2011) and, ii) FormiF/FormiR. The latter set was included because the ZBJ-Art primers have been reported to poorly detected ants and termites (Hamad et al. 2014) (Brandon-Mong et al. 2015). Thus we specifically designed this set to ensure detection of ants (Appendix Table S2), by first downloading all arthropod COI sequences from South Africa archived on GenBank in January 2016, then designing them by hand in Geneious 7.0.6 (Biomatters, New Zealand) to meet the following criteria: a) 20-22 bp length; b) produce a fragment of 150-210 bp (including primers); and c) C/G as the first two 3’ end bases (alignment provided as Appendix).

In addition, because robins have been seen eating berries (P. Lloyd, pers. comm.), we targeted the chloroplast rbcL gene using primers rbcL-h1aF/rbcL-h2aR (Poinar et al. 1998). The ZBJ-ArtF1c/ZBJ-ArtR2c primers amplify a 211 bp region in the 5’ section of COI gene, while the FormiF/FormiR primers target a 168bp region in the 3’ of same gene. All primers were modified to include a unique 6 to 8 nucleotide sequence (nucleotide multiplex identifiers; MIDs) on the 5’ end to allow individual identification following (Binladen et al. 2007). MIDs were designed with a custom script using R (R version 3.2.1): we generated random sequences of 6-8 nucleotides, where the probability of each base per site was 0.25, and selected those with pairwise differences >50% as our MIDs.

Prior to the metabarcoding PCR amplifications, we compared the efficiency of ZBJ-ArtF1c/ZBJ-ArtR2c and FormiF/FormiR primers, and generated pilot data to help decide the clustering thresholds needed to obtain the most accurate taxonomic richness, by performing PCRs on an artificial positive control. This sample contained a mixture of DNA derived from four species covering a broad taxonomic breadth (1 Coleoptera: Carabidae, 1 Diptera: Sarcophagida, 2 Hymenoptera: *Tetramorium sp* and *Camponotus fulvopilosus*).

PCR amplifications (25 µl) contained 1-3 µl of template DNA, 25 nM MgCl_2_ (stock 25nM), PCR buffer (stock 1x), 2 nM each dNTP, 0.6 nM each primer, 0.2U AmpliTaqGold DNA polymerase; thermal cycling conditions for arthropods amplifications (primers Formi and ZBJ) were: 95 °C for 5 min, followed by 33 cycles of 95 °C for 30 s, 48 °C for 30 s, and 72 °C for 30 s. Thermal cycling conditions for plant amplifications were 95 °C for 5 min, followed by 38 cycles of 95 °C for 30 s, 50.0 °C for 30 s and 72 °C for 30 s. A final extension step of 72 °C for 5 min was included in all reactions. The optimal number of PCR cycles was established using a real-time PCR assay in a subset of samples as the inflection point in the amplification curve and it represents a compromise to obtain sufficient prey DNA while reducing clonality and consequent bias in prey diversity. Real time PCR was performed in 20ul reaction containing 1-2 µl of template, 1x buffer, 2.5 nM MgCl2, 2 nM each dNTP, 0.5U Amplitaq Gold, 1 nM of each primer, and 1 µl SYBR Green/Rox mix (Invitrogen, USA). PCR amplifications were detected by gel electrophoresis on 2% agarose gel stained with Gelred. All PCR preparations used aerosol resistant filter tips, and were prepared in UV-sterilized laminar flow hoods. We note that arthropods and fecal samples were processed independently, and the same applied to summer and winter samples. This differential processing of samples was implemented as to avoid cross-contamination.

All PCR products were visualized on a gel to confirm amplification success, and then pooled at an approximately equimolar ratio into 11 pools (5 for summer and 6 for winter). We included the extraction blanks (from both fecal and arthropod pitfall extractions) in the pools, using 5 µl, despite no actual product visualized in agarose.

We removed unspecific fragments and primer dimer using Ampure XP SPRI beads, following a double-sided protocol. Shortly, by adjusting the ratio of beads to DNA (0.7x) we first captured the fragments larger than 300bp, keep the supernatant, which was in turn used to sequester fragments larger than 120bp (beads:DNA ratio = 1.8x). The concentration of the purified pools was determined in Qubit and efficiency of purification evaluated in and 2% agarose gel stained with Gelred.

The PCR pools were converted into sequencing libraries using the NEBNext 6070 blunt end library build kit following the manufacture’s protocol. Index PCRs were carried out in 25 µl reaction volumes, and each library was amplified in three replicates to maximize amplicon diversity. Each indexing PCR consisted of 2.5 µl of library, 1x Gold buffer, 2.5 U AmpliTaqGold DNA polymerase, 2.5 nM MgCl2 and 2 nM each dNTP, 0.2 µM forward indexed primer (InPE1.0), and 0.2 µM reverse index primer such that library contained a different reverse index. PCR was performed at 95°C for 5min, followed by 8 cycles at 95°C for 30 sec, 60°C for 30 sec, 72°C for 30 sec, and final extension at 72°C for 5 minutes. We combined the three replicates of each library and purified them using the QiaQuick columns (Qiagen, USA). To determine the quality and quantity (concentration) of our 11 libraries, we used the TapeStation System (Agilent Technologies, USA). We equalized the amplified library concentrations, then sequenced them on an Illumina MiSeq platform using PE250 chemistry and spiked with 30% of PhiX.

Metabarcoding amplifications were performed in triplicate in order to enable us to: i) discard potential errors (i.e. remove sequences only present in one PCR), and ii) optimize diversity (i.e. keep sequences only if they were present in at least two PCRs). All PCRs included one negative control to check for contamination and one positive control (DNA from an ant or plant species not present in southern Africa) to ascertain amplification. PCR products were incorporated into sequencing libraries (details in Appendix) for generation of metabarcoding data using an Illumina MiSeq platform with PE250 chemistry.

### 2.4 Sequence processing and taxonomic assignment

Post sequencing, the raw reads were processed with TRIMMOMATIC-0.36 (Bolger et al. 2014) to first remove adapters (ILLUMINACLIP module), then bases with low quality. The quality of the surviving reads (mean of surviving reads across pools: 93%) was checked with FASTQC (http://www.bioinformatics.babraham.ac.uk/projects/fastqc/) and then the overlapping paired-end reads were merged into contigs using PEAR (Hebert et al. 2003). These contigs were then demultiplexed and filtered with DAMe (Zepeda-Mendoza et al. 2016) to only retain fragments that matched the following criteria: i) present in at least two PCR replicates, ii) conform to the expected size after primer trimming, and iii) present in at least 25 copies (abundance cut-off), thus removing low abundance contigs that could be potential artefacts.

Clustering of sequences (within sample) into molecular operational taxonomic units (MOTUs) was carried out with USEARCH (Edgar 2010) and followed the 3% sequence divergence criteria. This clustering criteria was selected taking into account our empirical distribution of pairwise percentage of identity between arthropod sequences (Appendix Fig. 1S) as well as the reported variability in insect populations (Hebert et al. 2003). The MOTUs were aligned to the National Center for Biotechnology Information nucleotide sequence database (NCBI; website: http://www.ncbi.nlm.nih.gov/; accessed July 2017) the software Basic Local Alignment Search Tool (BLAST 2.2.6+; (Camacho et al. 2009)). The high-scoring hits obtained with BLASTn (e-value < 1e-30) were then parsed with MetaGenome Analyzer software (MEGAN v6.0; (Huson et al. 2007)) to assigning the MOTUS to their lowest common ancestor (LCA) in the NCBI taxonomic tree, as to meet the stringency criteria: min score = 130 for COI and 100 for rbcL, max expected = 0.01, min percent identity = 0, top percent = 10, min support percent = 0 (off), min support = 1, min complexity filter = 0.01. The MOTUs for which the LCA taxonomic assignment fell below the thresholds or for which a match was not found in the NCBI database was categorized as “unknown”. Not all MOTUs could be assigned to species level due to the incompleteness of the reference database for our study area. Therefore we present results identified to family-level. We contend this still provides a good grasp of the dietary components as well as the arthropod communities.

### 2.5 Statistical Analyses

All analyses were implemented in R v3.3.2 (R Development Core Team). We estimated Shannon-Wiener’s (H) and Simpon’s (1-D) indices to quantify both consumed and available prey diversity using *vegan* package (Oksanen 2017). Diversity indices were compared among sites using the non-parametric Kruskal-Wallis test, as the data was not normally distributed (Shapiro-Wilk test, p < 0.05). Changes in composition of prey were inspected using a non-metric multidimensional scaling (NMDS) with Bray-Curtis dissimilarity. To test whether region (*Coastal, Central, Inland;* proxy for primary productivity and energetic demands) is shaping diversity in available and consumed prey we used a Mantel test (Mantel 1967) with 999 random permutations as implemented in *vegan* (Oksanen 2017). Briefly, we compared the observed dissimilarity matrix (Bray-Curtis dissimilarity) against a conceptual matrix for regional effects, coded as zero for same region comparisons and one for different region comparisons. We highlight that we did not test for seasonal effects. Although we contend that seasonal variation in weather and a bird’s life-stage (breeding vs non-breeding) may have implications relating to available and consumed prey, respectively, and despite our effort to catch birds, the non-balanced sample sizes hinder a formal analysis.

Trophic niche breadth was estimated using a standardized Levin’s index (Bs; (Levins 1968). Standardized Levin’s index (Bs) was estimated as: Bs=(1/∑ p*i*^2^)-1/n-1, where p*i* is the proportion of individuals consuming arthropod family *i* and n is the total number of families available. This index measures the uniformity of distribution of individuals (birds) among resource states (arthropod families in each region) and it is closer to zero when diets are dominated by few items (specialist) and closer to one for a generalist diet. We created Bs 95% confidence intervals for each region by bootstrapping with 10,000 replicates. In addition, to test whether the observed Bs values were significantly different from a scenario of “no regional difference” we used a permutational approach. All occurrences (p*i*) were pooled regardless of their regional origin, then we randomly sampled *n* arthropod families (from the observed range: 37 – 45) and finally estimated Bs_random. This procedure was repeated 9999 times to generate the null distribution. The estimated probability of obtaining a result that exceeded the observed value under the null hypotheses of “no region difference” was estimated as p = [(number Bs_random > Bs_observed)/total number randomizations].

For the most common arthropod MOTUs in the diet of robins, we tested whether selectivity of prey changed among regions, by calculating Ivlev’s electivity index (Ivlev 1961) for each region as implemented in *selectapref* package. This index varies from “-1” (avoidance of prey) to “+1” (total reliance on prey), and “0” indicates prey selection is proportional to availability.

### 2.6 Ethics and Permits

Capture permits were issued by Cape Nature (0056-AAA008-00057) and the Department of Environment and Nature Conservation (1611/2015) in South Africa. Ethical approval was obtained from the Animal Research Ethics Committee at Nelson Mandela University (A15-SCI-ZOO-005).

## 3. RESULTS

### 3.1 Arthropod availability

Our survey of potential Arthropoda prey availability across the gradient revealed 519 MOTUs, with the majority (83%) identified to class Insecta (Appendix Table 3S). Coleoptera, Diptera, Hymenoptera and Lepidoptera were the most abundant orders (Fig. 2). The most diverse community was found in the *Central* region, the area with largest annual primary productivity (192 MOTUS, 45 Families, Shannon-Wiener’s H = 3.44, Simpson’s 1-D = 0.95; Table 1), whereas the in region with lowest NDVI and highest temperature amplitude, *Inland,* the arthropod community was least diverse (157 MOTUs, 42 Families, Shannon-Wiener’s H = 3.30, Simpson’s 1-D = 0.94; Table 1). However, when these observations were tested statistically, we were unable to reject the null hypothesis that there is no regional effect on arthropod community (Mantel_region_, p = 0.430).

**Table 1.** Available arthropod diversity and dietary diversity (Arthropod and Spermatophyta: seed plants) among populations of Karoo scrub-robin. T_amp_: annual temperature range. In parenthesis we denote relative arthropod consumption, i.e. % of prey consumed in relation to prey available.

|  | **Coastal** | **Central** | **Inland** |
| --- | --- | --- | --- |
|  | NDVI: 0.43 | NDVI: 0.50 | NDVI: 0.30 |
| | $T_{amp}$: 21.5 °C | $T_{amp}$: 27.7 °C | $T_{amp}$: 32.1 °C |
| Available Arthropod MOTUs | 170 | 192 | 157 |
| Available Families | 37 | 45 | 44 |
| Shannon-Wiener's H | 3.109 | 3.422 | 3.300 |
| Simpson's 1-D | 0.930 | 0.953 | 0.943 |
| Consumed Arthropod MOTUs | 36 (21%) | 57 (30%) | 64 (40%) |
| Consumed Families | 20 (54%) | 25 (56%) | 28 (64%) |
| Shannon-Wiener's H | 2.880 | 2.706 | 2.947 |
| Simpson's 1-D | 0.936 | 0.878 | 0.906 |
| Consumed Spermatophyta MOTUs | 21 | 22 | 43 |
| Consumed Families | 12 | 14 | 21 |
| Shannon-Wiener's H | 3.404 | 3.684 | 3.727 |

**Figure 2:**
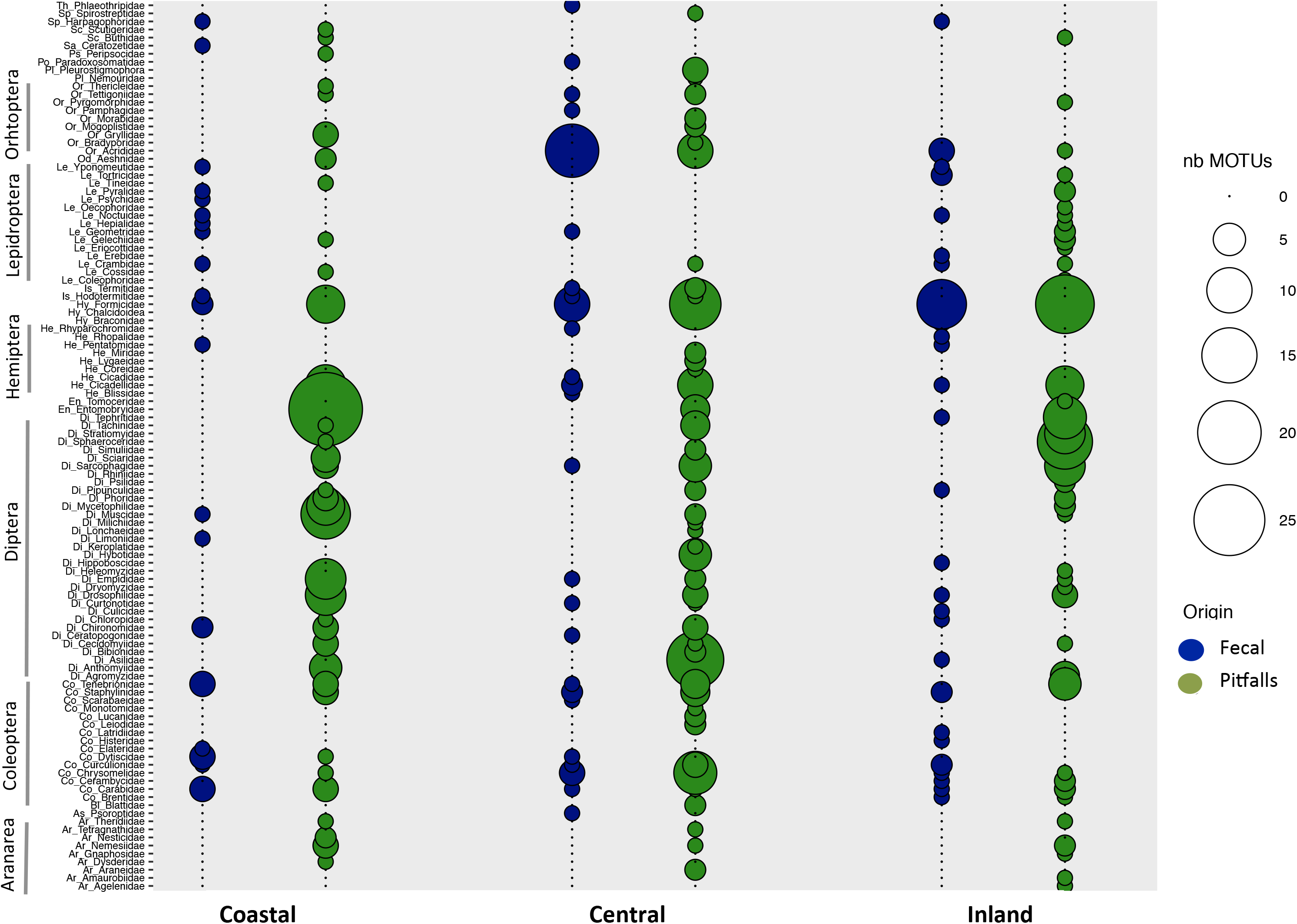
Diversity of Arthropod MOTUs per Family in pitfalls and fecal samples at three study locations: *Coastal*, *Central* and *Inland*. Bubble size is proportional to the number of MOTUs found in each family. Arthropod Orders are annotated with initials as a prefix to Family names (e.g.: Ar_Agelenidae): Ar: Araneae, As: Astigmata, Di: Diptera, En: Entomobryomorpha, He: Hemiptera, Hy: Hymenoptera, Is: Isopetra, Od: Odonata, Or: Orthtoptera, Pl: Pleurostigmophora, Po: Polydesmida, Ps: Psocoptera, Sc: Scutigemorpha, Sp: Spiristrepida, Th: Thysanoptera,

### 3.2 Arthropods and plants consumed

After data filtering criteria, 82% of fecal samples provided arthropod MOTUs (170). Insecta was the richest Arthropoda class (127 MOTUs) with the orders Coleoptera, Hymenoptera, Lepidoptera and Orthoptera being the most abundant (Fig. 2, Appendix Table 3S). Although the Karoo scrub-robin is viewed as insectivorous, we detected plant DNA in 94% of the fecal samples. Overall, 86 MOTUs were assigned to Spermatophyta (seed Plants) with Solanales and Aspargales being the most abundant (Appendix Table 4S).

Our data showed no evidence that dietary composition was associated with region whether considering only arthropod prey (Shannon-Wiener’s H: Kruskal-Wallis *Χ*^2^ = 1.855, df = 2, p = 0.396; Simpson’s 1-D: Kruskal-Wallis *Χ*^2^ = 1.885, df = 2, p = 0.390; Mantel_region_, p = 0.072; Fig. 3) or combing arthropod prey and plant items (Shannon-Wiener’s H: Kruskal-Wallis *Χ*^2^ = 0.745, df = 2, p = 0.689; Simpson’s 1-D: Kruskal-Wallis *Χ*^2^ = 0.422, df = 2, p = 0.810; Mantel_region_ p = 0.433; Fig. 3).

**Figure 3.**
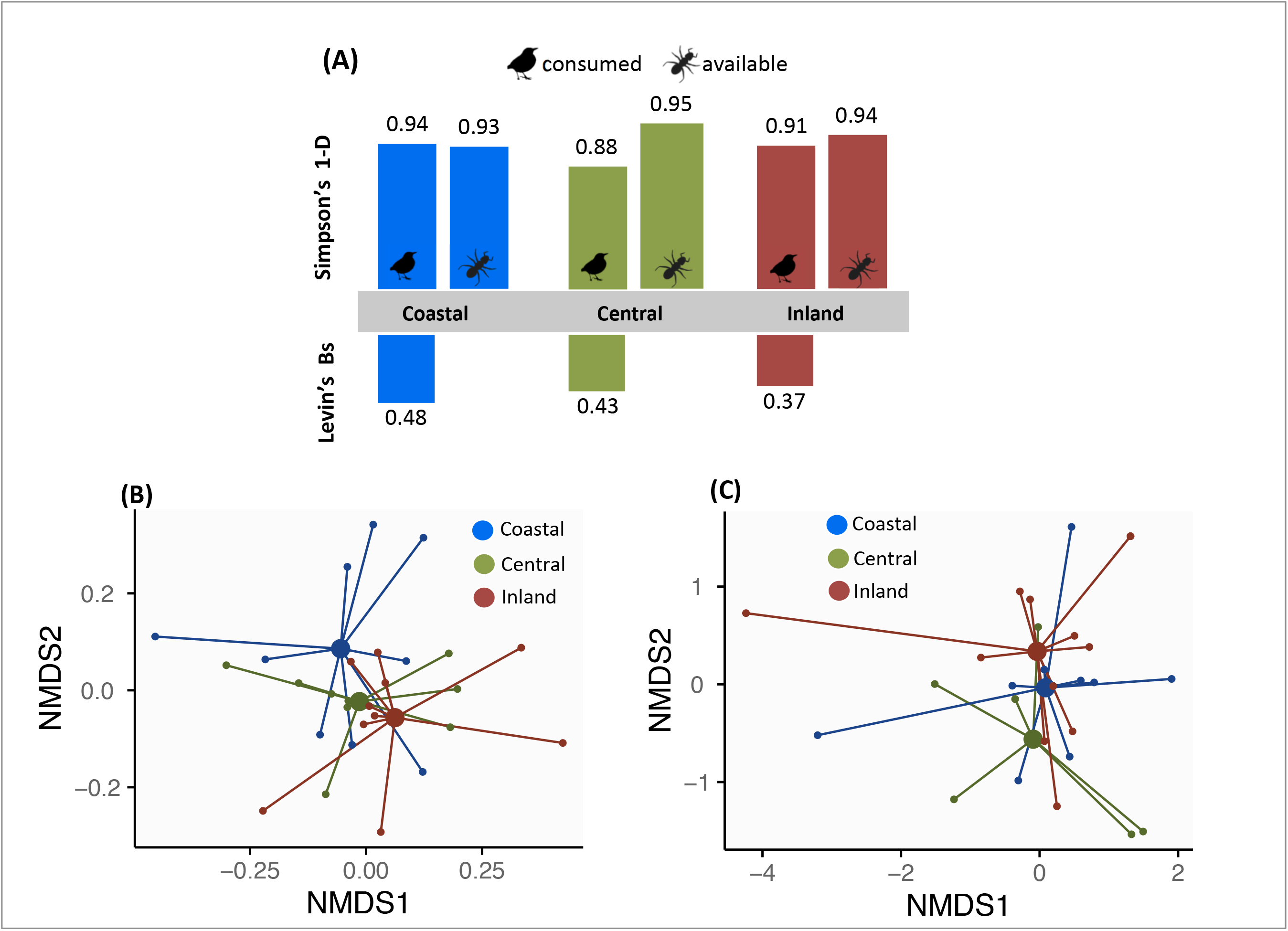
Diversity in arthropods potentially available to, and consumed by, the Karoo-scrub robin. (A) Simpson’s 1-D diversity index and Levin’s Bs niche breadth (Bs_*Coastals*_ 95% CI = 0.328 – 0.667, Bs_*centrall*_ 95% CI = 0.285 – 0.659, Bs_*Inland*_ 95% CI = 0.241 – 0.677). (B) NMDS for arthropod prey and (C) NMDS for plant items consumed; STRESS_arthropods_ = 0.17, STRESS_plants_ = 0.14.

### 3.3 Trophic niche and prey selection

When accounting for available arthropod prey, we found that birds living in the *Inland* region-which was least productive and exerted high energetic demands on the robins – appeared to have the narrowest dietary niche breadth (Bs = 0.372, 95% CI = 0.241 – 0.677, Fig. 3A). This contrasts with the *Coastal* population, whose individuals presented the widest niche (Bs = 0.479, 95% CI = 0.328 – 0.667; Fig. 3A). However, again when statistically tested, we found that the observed Bs values for each of the three study regions was not significantly different from the values expected under the null hypothesis of no regional difference (mean Bs_random = 0.445, 95% CI = 0.247 −0.592, p_*Coastal*_= 0.658, p_*Central*_ = 0.568, p_*Inland*_ = 0.738; Appendix Fig. 2S).

An electivity analysis (that measures utilization of food items) restricted to the five most abundant potential prey items (Coleoptera, Diptera, Hymenoptera, Lepidoptera and Orthoptera), showed that robins from all regions positively selected for Hymenoptera:Formicidae (Ivlev’s electivity > 0; Fig. 4). Electivity was, however, largest in the *Inland* population, with 55% of sampled birds consuming ants. Although in *Central* region relatively more Coleoptera families were potentially available as prey (Appendix Fig. 3S), robins did not positively selected on this prey type (Fig. 4). We also found that Dipterans, another widely available potential prey source, were negatively selected on in all regions (Ivlev’s electivity < 0; Fig. 4)

**Figure 4.**
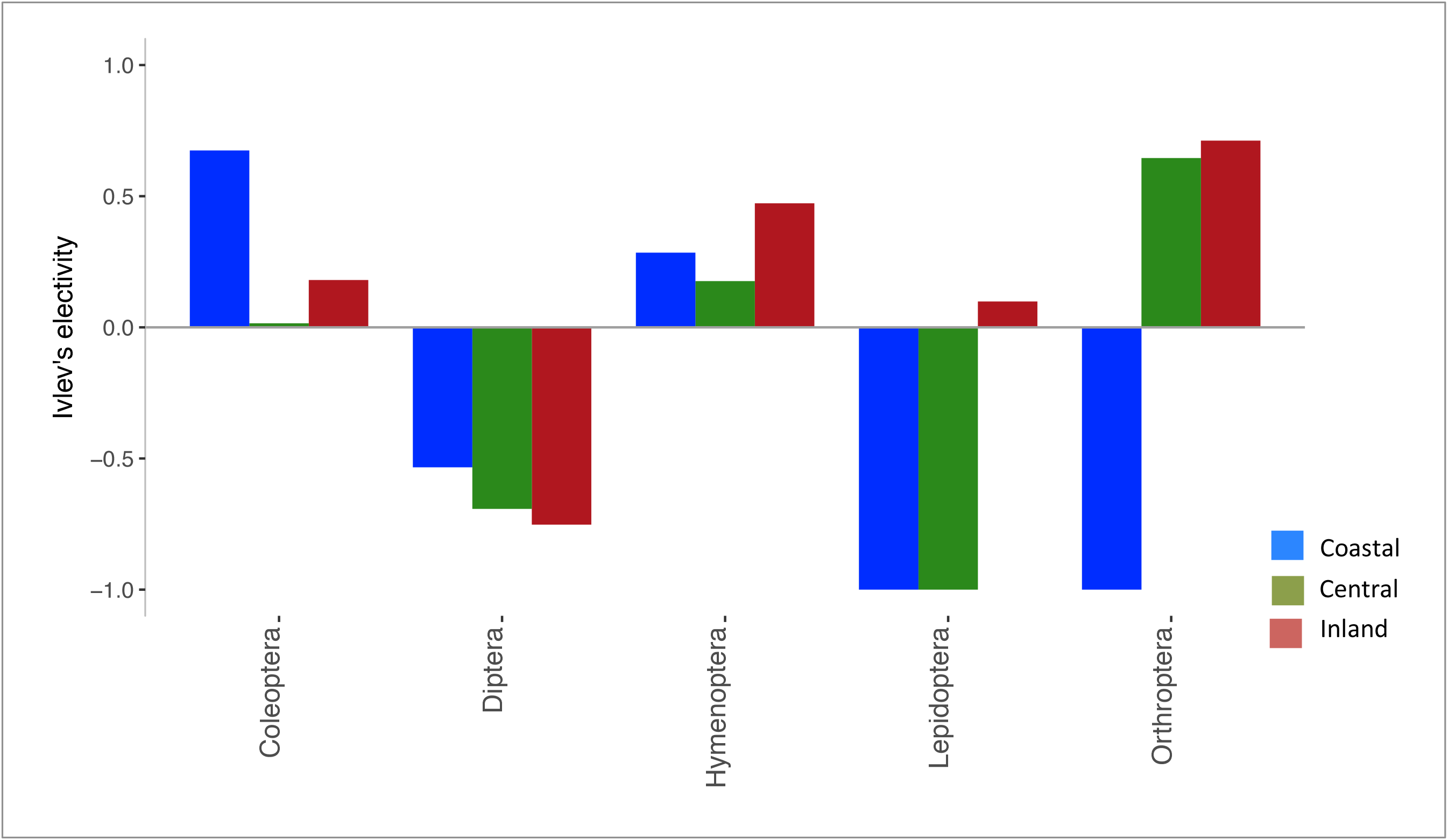
Ivlev’s electivity index for the main arthropod orders consumed: Coleoptera (Co) Diptera (Di), Hymenoptera (Hy), Lepidoptera (Le) and Orthoptera (Or). Positive values indicate prey selection whereas negative values indicate prey avoidance.

## 4. DISCUSSION

Species living in spatially and temporally heterogeneous environments must adjust their behavior and physiology in order to persist. Dietary plasticity may thus be an advantageous trait in energy-depauperated ecosystems as switching from alternative resources allows them to deal with environmental change (e.g.: (Varner and Dearing 2014). Here, we investigated dietary patterns among conspecific Karoo scrub-robin populations living in regions with different productivity and thermoregulatory demands. Overall our findings revealed that: i) although Karoo scrub-robins are generalist insectivores, some plant items seem to be included in its diet; ii) dietary composition was not associated with region; iii) when accounting for available arthropod prey the dietary niche breadth of each population was not different from a scenario of no regional effect (i.e., similar productivity and thermoregulatory demands); iv) Formicidae (ants) were positively selected in the three populations, yet *Inland* robins showed the largest electivity.

### Dietary niche breadth

The concomitant assessment of available and consumed prey in conspecific populations revealed that the foraging behavior of robins is not simply based on encounter rate: Coleopterans, Hymenopterans and Orthopeterans were preferred prey. While coleopterans represent less than 16% of available prey (7%, 16% and 11% of MOUTUs in *Coastal, Central* and *Inland*, respectively) they were clearly preferentially preyed upon (representing 33% of total dietary MOTUs in the *Coastal* region, 23% in the *Central* region and 20% in the *Inland* region). A similar pattern of preference was found for Hymenopterans in the *Central* and *Inland* regions: Hymenopetans were consumed at a higher rate than their relative availability (comprising 11% and 19% of the dietary MOTUs, *vs* 7% and 11% of available MOTUs). The exception was found in the area with intermediate primary productivity and temperature amplitude – the *Coastal* region – where Hymenopetans were only 4% of available prey and 5% of consumed MOTUs. In contrast, Diptera and Lepidoptera were consumed at a rate inversely proportional to their presence, i.e. they were avoided by the birds. These results are in accordance to prey descriptions based on Karoo scrub-robins stomach contents (invasive method) sampled throughout the year in a region East of our *Inland* site (Free State; (Oatley 1970)). Specifically, Oatley (1970) revealed that ants (Hymenoptera) and beetles (Coleoptera) were present in 55% and 14% of the stomachs, respectively. We also noted that while terrestrial arthropods are the bulk of robin’s prey, they appear to supplement their diets with plant matter (94% of samples yielded plant DNA). Although, this finding could in theory simply indicate that robins are plant secondary consumer through herbivore arthropods, we contend the latter is unlikely given both the anticipated degradation to the plant DNA that would be incurred through two rounds of ingestion and degradation, and perhaps more importantly, two additional lines of evidence. Firstly, there are unpublished observations (P. Lloyd, pers. comm.) of birds foraging on berries and plant shoots, and secondly, seeds and berries were detected in the stomachs contents of robins (Oatley 1970). Thus together, it suggests that that plant matter is not simply a serendipitous food item, but may in fact be selected for.

Our study sites are embedded within the Southern African semi- and arid-regions, where seasonal pulses of rainfall are known to determine arthropod densities (Dean and Milton 1999) and subsequently food supplies for insectivores (Lloyd 1999). Surprisingly, despite the differences in primary productivity among our study sites, the dietary niche breadth (Bs) among populations was not different from a scenario of homogeneous environment (i.e., no difference in productivity or thermal demands). This plasticity in diet may be favorable in environments where prey is patchily distributed in space and time, such as the Southern African semi- and arid-regions (Dean and Milton 1999). Moreover, this finding corroborates the expectations of Optimal Foraging Theory that animals can only afford to be specialist (small niche breadth) where resources are readily available (Stephens and Krebs 1986).

### Life in arid environments

Life in arid environments is harsh for endotherms in general, and in particular for ground-feeding insectivore birds as they need to fuel their typically high avian metabolic rates in energy-depauperated ecosystems. We predicted the robins living in the most demanding regions (*Inland*) would exhibit the most generalist feeding strategy, under the rationale that as this was the most energy-depauperated ecosystem and energetically demanding environment, their diet should become more generalist as ignoring potential sources of energy is not an option (Levins and MacArthur 1969; Schoener 1971). Specifically, we expected robins living in the *Inland* region to prey on the greatest diversity of ground arthropods, so as to obtain sufficient energy to fuel the thermogenic demands of winter (burn energy to produce heat as a means to warm-up (thermogenesis; (Hothola 2004) as well as actively cool down in summer through evaporative cooling (Williams and Tieleman 2005). In contrast however, we found that the *Inland* robins prey on an array of arthropods that is similar to robins living in the less demanding region (*Coastal*). We also noted they exhibit a preference for ants (Formicidae), a finding that lends support to the notion that dependable food sources such as ants are critical for arid endemic birds (Dean and Milton 2017), yielding the necessary energy, nutrients and water (robins do not drink) to overcome the challenges that the environment exerts.

We also hypothesize that supplementation of their diet with plant items such as berries (the most common plant in diet, Solanaceae, produces berries) may provide the carbohydrates and hence calories required to cover their general energetic budget, when sufficient animal protein and fat is not available. Furthermore beside sugars, fruits would also provide extra water, a critical component when dealing with hot temperatures: birds loose water to cool down and avoid hyperthermia in the hot days. Although we did not test for seasonal differences, it is worth noting that plant DNA was mostly found in samples collected in winter. We hypothesize this may be related to high energy demands to deal with cold as well as short time-window to actively search for arthropod prey on the ground (shorter periods of day light).

### eDNA metabarcoding in trophic ecology

Investigating foraging strategies by applying molecular methods to environmental and non-invasive samples is increasingly facilitating the study of highly-mobile and evasive animals (e.g.: (Jedlicka et al. 2013) (Mallott et al. 2014) (Jedlicka et al. 2017). Nevertheless, there are several caveats underlying inferences of the dietary components and arthropod communities based on DNA recovered from fecal and environmental samples: i) data is typically qualitative allowing only presence/absence inferences, ii) primer specificity may bias detection of some taxa, iii) the differential digestibility of hard- *vs* soft-bodied prey may bias results, iv) incompleteness of reference databases can hinder species-level taxonomic assignment, v) samples, either fecal or environmental, represent a snapshot of the dietary diversity available and preyed on, and iv) inferences of herbivory may be confounded if animals are actively consuming herbivorous arthropods – secondary consumption. Nevertheless, we contend the dietary patterns obtained in this study are robust and provide a first grasp on how arid endemic birds may be ensuring sufficient energy yields in a rather challenging environment. Thus we hope our findings provide a stepping-stone for further research into the role of plasticity in foraging strategies. Moreover, studies like this one are both relevant for evolutionary biologists because describing the factors that lead to dietary variation in natural populations is fundamental for understanding local adaptation and speciation (e.g.: (Grant et al. 2008; Svanbäck and Schluter 2012), as well as for conservation biologists because the on-going global change is expected to increase aridity and consequently alter food-webs, which in turn affect avian communities (e.g.: (Iknayan and Beissinger 2018).

## Supporting information

Supp. Methods and Results

## Acknowledgments

We thank the Southern African authorities for providing permits, the Danish National High-Throughput DNA Sequencing Centre for generating sequence data, Nick Pattinson for his help in the field, Lara Puetz and Sarah Mak for useful lab hints, Shyam Gopalakrishnan for assistance with bioinformatics, Rute Fonseca for valuable discussions and Penn Lloyd for sharing is observations about the foraging behavior of Karoo scrub-robins. This study was funded by a Marie Sklodowska-Curie Individual fellowship to AMR (Grant 655150S-BARREN).

## AUTHOR CONTRIBUTIONS

AMR and MTPG conceived the study. AMR and BS collected samples. AMR performed molecular work and conducted analyses. AMR, BS and MTPG wrote the paper.

## DATA ARCHIVING

Data reported are available as Appendix.

1. Fasta file with Formi primers aligned against Formicidae.
2. Fasta files with available and consumed MOTUs obtained with different primer sets.

