## Supplementary material for "31° South: Dietary niche of an arid-zone endemic passerine": Supp. Methods and Results

**Supplementary Material for manuscript "31° South: Dietary niche of an arid-zone endemic passerine" by Ribeiro et al.**

<u>**SUPLEMENTARY TABLES**</u>

**Table 1S**: Large arthropods excluded from 'arthropod soup' to avoid biasing DNA extraction, but added in the final dataset after morphological identification.

| Coastal | Central | Inland |
|---|---|---|
| **Tenebrionidae:** Psammodes sp | **Tenebrionidae:** Psammodes sp | **Tenebrionidae:** Psammodes sp |
| **Carabidae:** Thermophilum *sp* Anthia maxillosa | **Bradyporidae:** Acanthoplus discoidalis | **Carabidae:** Thermophilum decemguttatum |
| | | **Buthidae:** *Parabuthus sp* |

**Table 2S.** Primer sequences (5′–3′) used to amplify arthropods and plant DNA from Karoo Scrub-robin fecal samples.

| Primer name | Sequence 5'-3' | Amplicon size (bp) | Target COI barcode (bp) | Reference |
|---|---|---|---|---|
| ZBJ-ArtF1c | AGATATTGGAACWTTATATTTTATTTTTGG | 211 | | Zeale et al 2011 |
| ZBJ-ArtR2c | WACTAATCAATTWCCAAATCCTCC | | | |
| Formi_F | GGAATRTCWTCAATYYTMGG | 167 | 437 | This study |
| Formi_R | AGTATGGTAATDGCHCCDGC | | 605 | |
| rbcL h1aF | CAGCATTCCGAGTAACTCCTC | 136 | | Poinar et al 1998 |
| rbcL h2aR | CGTCCTTTGTAACGATCAAG | | | |

**Table 3S.** Details about molecularly detected OTUs in pitfalls and fecal samples, its taxonomic assignment and diversity. The taxonomic assignment performed with MEGAN v6 includes MOTUs assigned to the Family node and all the descendent nodes in the taxonomic tree. Some large Arthropods from Pitfalls were identified to Family-level by morphological characters and added to the final dataset for available prey (Metagenomics+Morphology).

| | | Available Prey (pitfalls) | | | Consumed Prey (fecal) | | |
|---|---|---|---|---|---|---|---|
| | | **Coastal** | **Central** | **Inland** | **Coastal** | **Central** | **Inland** |
| **Metabarcode** | Total MOTUs | 213 | 346 | 278 | 46 | 69 | 84 |
| | MOTUs assigned | 170 | 198 | 157 | 38 | 61 | 69 |
| | Arthropoda | 170 | 192 | 157 | 36 | 57 | 64 |
| | Arachnida | 8 | 10 | 9 | 1 | 2 | 0 |
| | Chilopoda | 1 | 4 | 0 | 0 | 0 | 0 |
| | Diplopoda | 0 | 1 | 0 | 0 | 1 | 1 |
| | Maxilopoda | 1 | 2 | 0 | 0 | 0 | 0 |
| | Collembola | 28 | 8 | 4 | 0 | 0 | 0 |
| | Insecta | 131 | 154 | 144 | 35 | 52 | 63 |
| | Blattodea | 0 | 10 | 0 | 1 | 2 | 2 |
| | Coleoptera | 13 | 30 | 17 | 12 | 13 | 13 |
| | Diptera | 70 | 61 | 73 | 6 | 4 | 9 |
| | Hemiptera | 8 | 11 | 7 | 1 | 5 | 3 |
| | Hymenoptera | 7 | 13 | 17 | 2 | 6 | 12 |
| | Lepidoptera | 17 | 4 | 25 | 6 | 3 | 12 |
| | Odonata | 2 | 0 | 0 | 0 | 0 | 0 |
| | Orthoptera | 5 | 14 | 2 | 0 | 14 | 3 |
| | Plecoptera | 0 | 1 | 0 | 0 | 0 | 0 |
| | Psocoptera | 3 | 2 | 0 | 0 | 0 | 2 |
| | Thysanoptera | 0 | 0 | 0 | 0 | 1 | 0 |
| | Number Families | 37 | 45 | 42 | 20 | 25 | 28 |
| | MOTUs Family assigned | 135 | 130 | 119 | 27 | 47 | 45 |
| **Metabarcode + Morphology** | MOTUs Family assigned | 138 | 133 | 122 | 27 | 47 | 45 |

**Table 4S.** Plant MOTUs detected in fecal samples: details about taxonomic assignment and diversity indices.

| | **Consumed (fecal)** | | |
|---|---|---|---|
| | **Coastal** | **Central** | **Inland** |
| MOTUs | 28 | 53 | 81 |
| MOTUs assigned | 21 | 35 | 50 |
| Spermatophyta (Seed Plants) | 21 | 22 | 43 |
| Alismatales | 1 | 0 | 0 |
| Apiales | 0 | 0 | 2 |
| Asparagales | 5 | 2 | 1 |
| Asterales | 0 | 2 | 0 |
| Brassicales | 0 | 0 | 1 |
| Caryophyllales | 2 | 0 | 5 |
| Cornales | 0 | 0 | 1 |
| Cupressales | 1 | 0 | 0 |
| Ericales | 0 | 0 | 1 |
| Fabales | 2 | 1 | 3 |
| Fagales | 2 | 0 | 1 |
| Gentianales | 0 | 2 | 1 |
| Lamiales | 1 | 1 | 3 |
| Malvales | 0 | 0 | 1 |
| Oxalidales | 1 | 3 | 0 |
| Pandanales | 0 | 0 | 1 |
| Poales | 0 | 2 | 1 |
| Santales | 0 | 2 | 0 |
| Sapindales | 0 | 0 | 1 |
| Saxifragales | 0 | 0 | 3 |
| Solanales | 5 | 2 | 13 |
| Zingiberales | 0 | 1 | 0 |
| Zygophyllales | 1 | 2 | 2 |
| Number families detected | 12 | 14 | 21 |
| MOTUS assigned Family level | 21 | 20 | 39 |

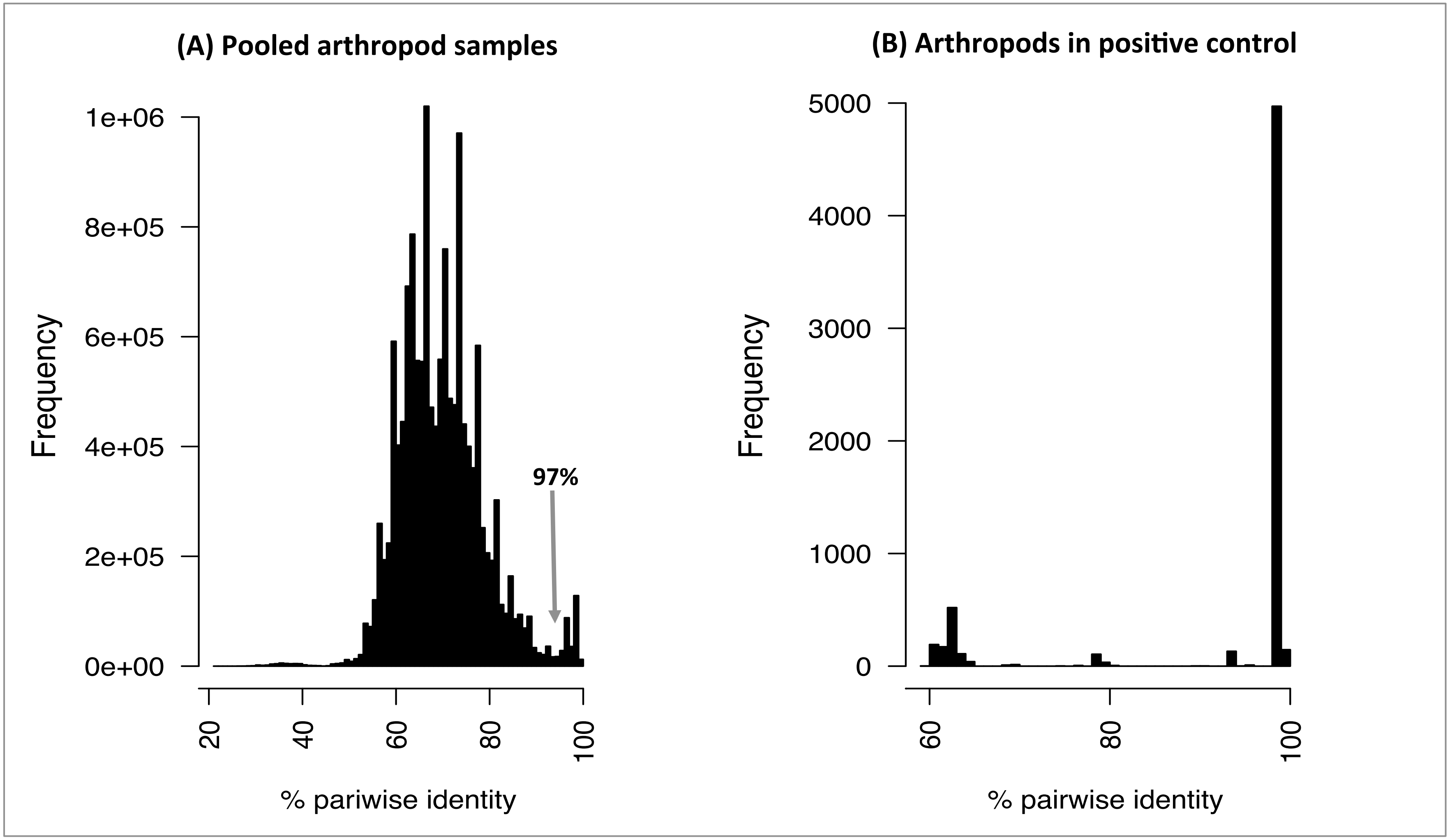


**Figure 1S**. Distribution of pairwise percentage of identity among COI mini-barcodes amplified with Formi primers for (A) all arthropod samples (pitfalls) and (B) positive control with four different species. The sequences were aligned with MAFFT (Katoh 2002) and show a break at 96-97%.

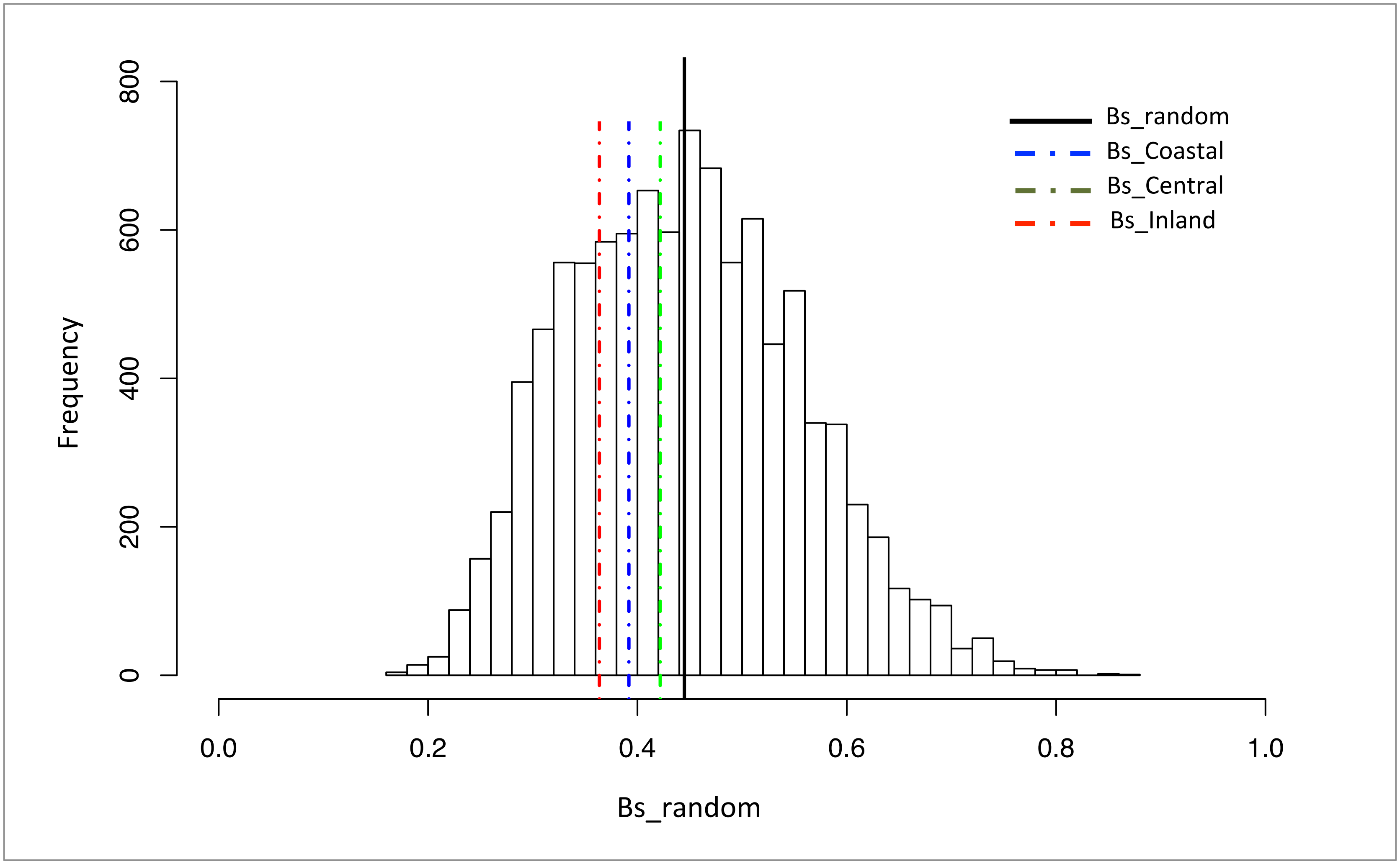


**Figure 2S**. Distribution of Levin's niche breadth (BS) under a scenario of "no region differences" obtained by permutation (10,000). Mean value of simulated BS (BS_random) and observed Bs values for each region are depicted as vertical lines across the histogram. Bs observed in each region was not significantly different from simulated scenario (p> 0.1).

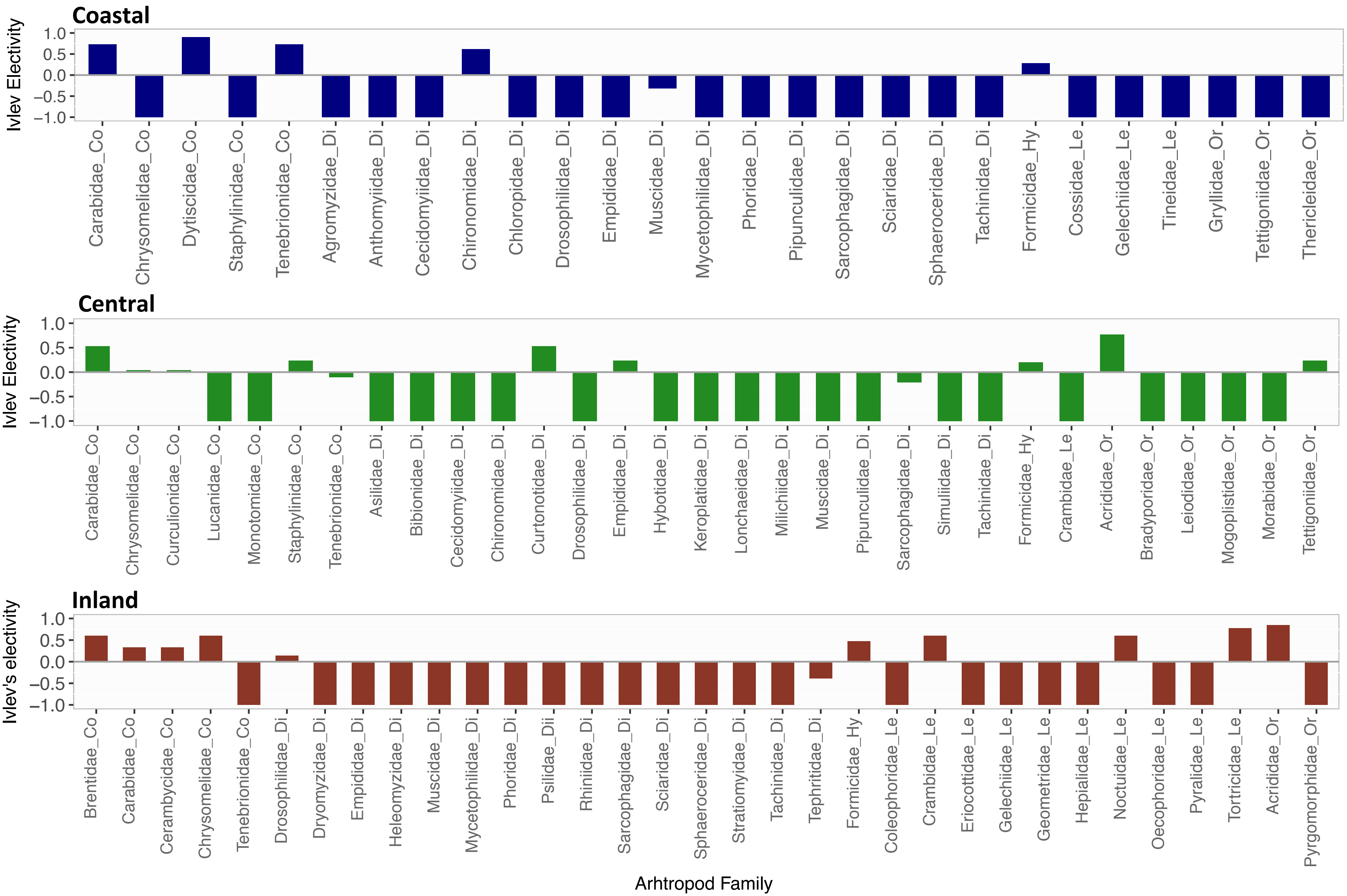


**Figure 3S** Ivlev's electivity index for the main arthropod orders consumed: Coleoptera (C) Diptera (D), Hymenoptera (H), Lepidoptera (L) and Orthoptera (O). Positive values indicate prey selection whereas negative values indicate prey avoidance.
